# Seasonal polyphenism of wing colors and its influence on sulphur butterfly diversification

**DOI:** 10.1101/2022.08.10.503521

**Authors:** Jennifer Fenner, Vincent Ficarrotta, Alexandra Colombara, Heather Smith, Kymberlee Evans, Ryan Range, Brian A. Counterman

## Abstract

Seasonal variation of color patterns on butterfly wings are iconic examples of developmentally plastic traits that can influence adaptation and speciation. Yet, there are few examples of such seasonal polyphenisms that have characterized the environmental cues, ecological consequences, or genetic mechanisms involved in generating the plastic variation of wing color. Further, there is a lack of support that such plasticity may impact the adaptive diversification of butterfly wing patterns. Here, we report a case of seasonal polyphenism in pigment and structurally-based color patterns of *Zerene cesonia* that are strikingly similar to the color pattern divergence seen on the wings of sulphur butterflies. We show that (i) coordinated changes in temperature and photoperiod drive the plasticity, (ii) the plastic color changes impact how fast the butterflies can warm, (iii) identify *spalt* as likely be involved in the genetic coupling of the pigment and structurally-based color plastic response. We further show that this plastic wing changes phenocopy wing pattern divergence between *Zerene* species, as well as the color pattern differences known to be commonly involved in sexual selection and speciation across sulphur butterflies. Together, our results demonstrate that shared environmental cues and genetic basis for pigment and structural color plasticity may result in conditions that may have facilitated species diversification of sulphur butterflies.

## Introduction

Developmental plasticity, an organism’s ability to develop differently in response to its environment, is a major driver of phenotypic diversity. Seasonal polyphenisms found in a broad range of organisms are powerful examples of how the environment can impact developmental trajectories and trait variation (1, 2). Butterfly wing color patterns offer some of the most striking examples of seasonal polyphenisms (3). Importantly, many of these polyphenisms have been shown to be adaptive and influence mate preference (4–10). Thus, seasonal polyphenisms of butterfly wings offer the opportunity to investigate how developmental plasticity may provide the phenotypic variation to fuel adaptive diversification and species divergence.

Genetic variation in the plasticity of a trait is a major determinant in the ability of plasticity to influence evolutionary processes. Although natural and sexual selection cannot act on environmentally induced trait variation, they can act on heritable genetic variation that influences the plasticity of a trait. Waddington demonstrated this in his classic experiments of “genetic assimilation” in *Drosophila*, where he used artificial selection among laboratory lines to alter plastic responses in varying environmental conditions (11). This experiment was recently repeated in *Junonia* butterflies, where van der Burg et al (12) successfully demonstrated the genetic assimilation of plastic wing colors, and identified candidate genes underlying the changes in plasticity. This study demonstrates the clear potential for selection to act on the seasonal plasticity of butterfly wing color patterns and influence the genetic divergence of populations.

Here, we characterize seasonal polyphenism in wing color patterns of *Zerene cesonia* butterflies and examine the possibility that such plasticity could have influenced the diversification of sulphur (subfamily Coliadinae) butterflies. In the sister genus, *Colias* butterflies, seasonal polyphenism in wing pigmentation is adaptive. The darker morphs of colder months are able to warm more quickly than lighter colored ones (9), which increases their opportunities for mating and oviposition. In *Zerene*, we observed seasonal variation in not only pigmentation but also structurally-based ultraviolet iridescent (UVI) wing patterns. We confirmed the seasonal polyphenism of pigment and UVI wing patterns in the laboratory and demonstrated that pigment and UVI plasticity shared similar environmental cues. Similar to *Colias*, the darker colored pink morphs warmed faster, suggesting the plasticity may have a similar adaptive benefit in *Z. cesonia*. Interestingly, the homologous marginal and discal cell melanic spots that showed seasonal polyphenism in *Zerene* are prefigured by the expression of the transcription factor *spalt* in *Colias* butterflies (13). Our CRISPR edits (crispants) of *spalt* in *Zerene* showed increased pink pigmentation and UVI loss, revealing coordinated changes in pigment and UVI that recapitulate endogenous seasonal wing color plasticity. Finally, we tested if UVI patterns differed among closely related Coliadinae species in a fashion that is similar to the plastic variation in *Z. cesonia* and discovered that 50% of species pairs studied differed in UVI presence/absence across the wing, including *Z. cesonia* and its sister species *Z. eurydice*. This supports that plasticity can generate the same patterns of UVI variation known to influence mate choice and hybridization among Coliadinae butterflies (14–20). Collectively, our findings support that the environmental and genetic changes involved in the seasonal polyphenism of pigment and UVI wing color patterns have the potential to influence adaption and speciation in Coliadinae butterflies.

## Results

### Seasonal polyphenism of pink pigmentation and structural UVI on Z. cesonia wings

Pigment and UV coloration varied seasonally in collections from northeast Mississippi. From late Spring to early Fall, the butterflies are predominantly yellow with little or no pink coloration, and a UVI pattern is present on all male dorsal hindwings (Figure 1). In late Fall and early Spring, all butterflies exhibited pink pigmentation (aka “*rosa*” morph) on the ventral wing surfaces, and 60% of males lacked a UV pattern on the hindwings. An assessment of pink and UVI coloration across the wings revealed that despite more continuous variation in pink plastic responses (Figure 1), UVI presence /absence varied in a more switch-like fashion with any males that had >6% pink coloration on the ventral hindwing surface, always lacked a UVI pattern on the dorsal hindwing.

**Figure 1.**
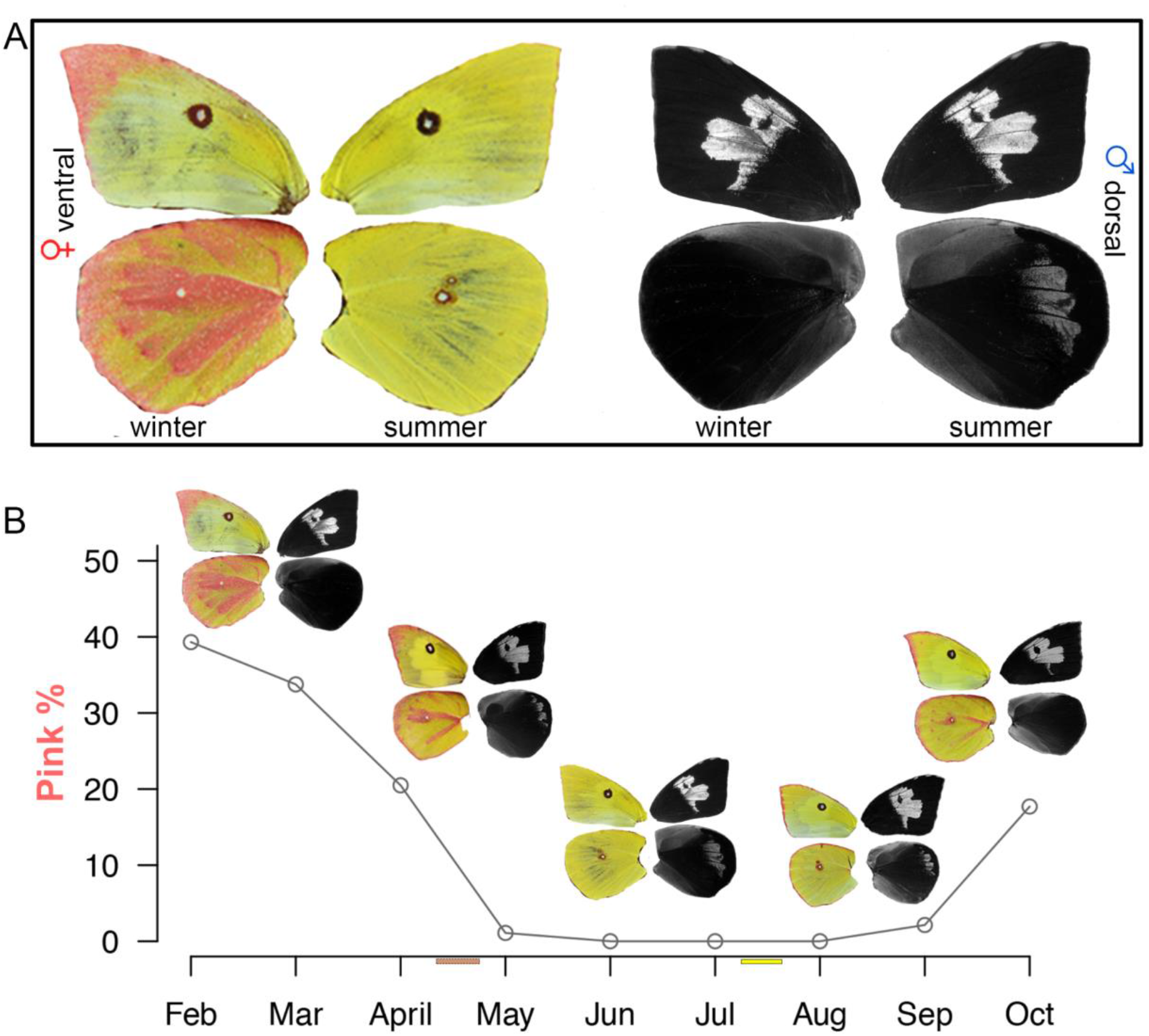
Pink and UVI wing color seasonal polyphenisms in *Z. cesonia*. **A**. Wild-caught *Z. cesonia* from winter and summer months (left visual; right UVI). **B**. Seasonal variation in pink pigmentation and UVI pattern in Mississippi population.

Pink pigmentation varied as a quantitative trait and increased during months with colder temperatures and shorter photoperiods. The average amount of pink coloration on the hindwing of an individual butterfly was on average 20% higher in the months of February and October (average maximum and minimum temperatures of 24°C and 5.3°C, respectively; approximately 12:12 photoperiods) than in the months of April and September (average maximum and minimum temperatures of 27.5°C and 15.5°C, respectively; approximately 14:10 photoperiods) (Figure 1B and 2B).

### Pigment and structural color plasticity share an environmental cue

Laboratory rearing experiments revealed that pigment and UV plasticity required changes in temperature and photoperiod (Figure 2). We used temperature and photoperiod conditions that reflected the environmental conditions in late-July (warm temperature; long photoperiod (W/LP)) and late April (cold temperature, short photoperiod (C/SP) to rear *Z. cesonia* in the laboratory (Figure 1B). Short photoperiod did not increase pink pigmentation or result in the loss of hindwing UV (Figure 2A, S1). Cooler temperatures resulted in a slight increase of pink pigmentation and a loss of hindwing UV for 25% of the males (Figure 2, S1). However, coordinated changes in both temperature and photoperiod did result in changes in both pigmentation and UVI. Reaction norm-like plots of pigmentation and UV show that by changing photoperiod and temperature in the lab we were able to recapitulate the plasticity seen in nature (Figure 2B). The only exception was that in the lab-reared W/LP conditions, the percent of hindwing with UVI was considerably greater than in wild-caught individuals. Collectively, these findings demonstrate that the plasticity of pigmentation and UVI share a similar environmental cue, requiring coordinated changes in both temperature and photoperiod.

**Figure 2.**
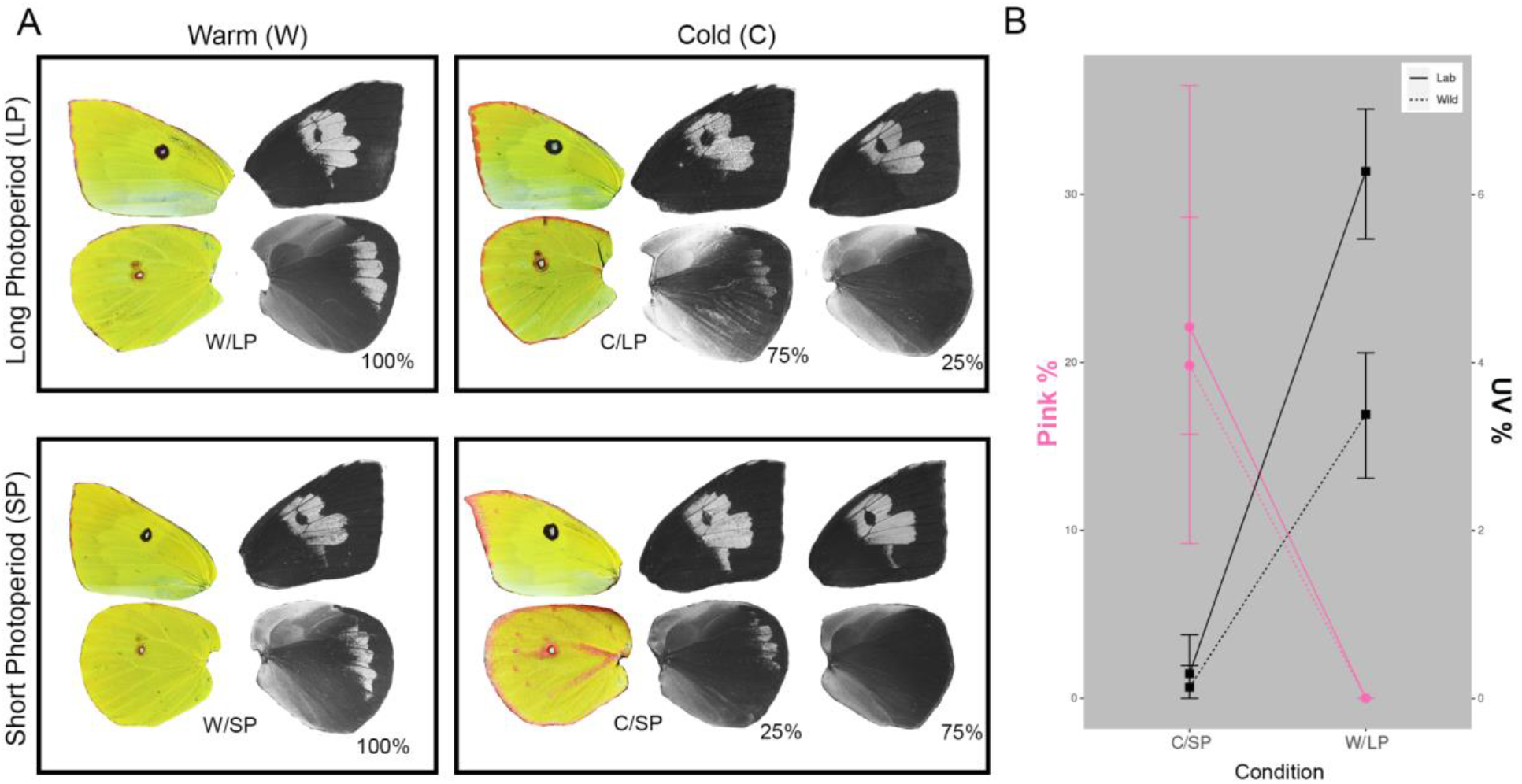
Pink and UVI plasticity are induced by changes in temperature and photoperiod. **A**. Both seasonal forms were successfully recreated in the laboratory through conditional rearing. Changes in photoperiod and temperature are needed to induce changes in pink and UVI. **B**. Reaction norm type plots of pink and UVI pattern changes in lab-reared and wild-caught individuals. Individuals for wild-caught were collected in two intervals denoted by horizontal pink and yellow bars in Figure 1B, when field conditions were most similar to C/SP and W/LP lab conditions, respectively.

### Pink butterfly wings have distinct scale types and warm faster

The wings of pink winter form butterflies in wild populations develop an assortment of pink, white, and yellow scales in narrow regions across the forewing and hindwing ventral surfaces (Figure S2-3). All three scale types show a high density of pterin pigment granules and share the typical ultrastructure of pterin wing scales. The scales of winter forms also have lamellar surfaces and ridge structures that are distinct from melanic scales from the same wing regions of summer form individuals. The pink scales are significantly taller and wider than yellow scales, and wider than melanic scales (Figure S2). The pink pigmentation results in darkly toned wing regions, particularly near lower margin spots on the hindwing. The result is that winter forms can have as much as 40% of their hindwing ventral surface covered by large, pink-pigmented scales (Figure 1).

**Figure 3.**
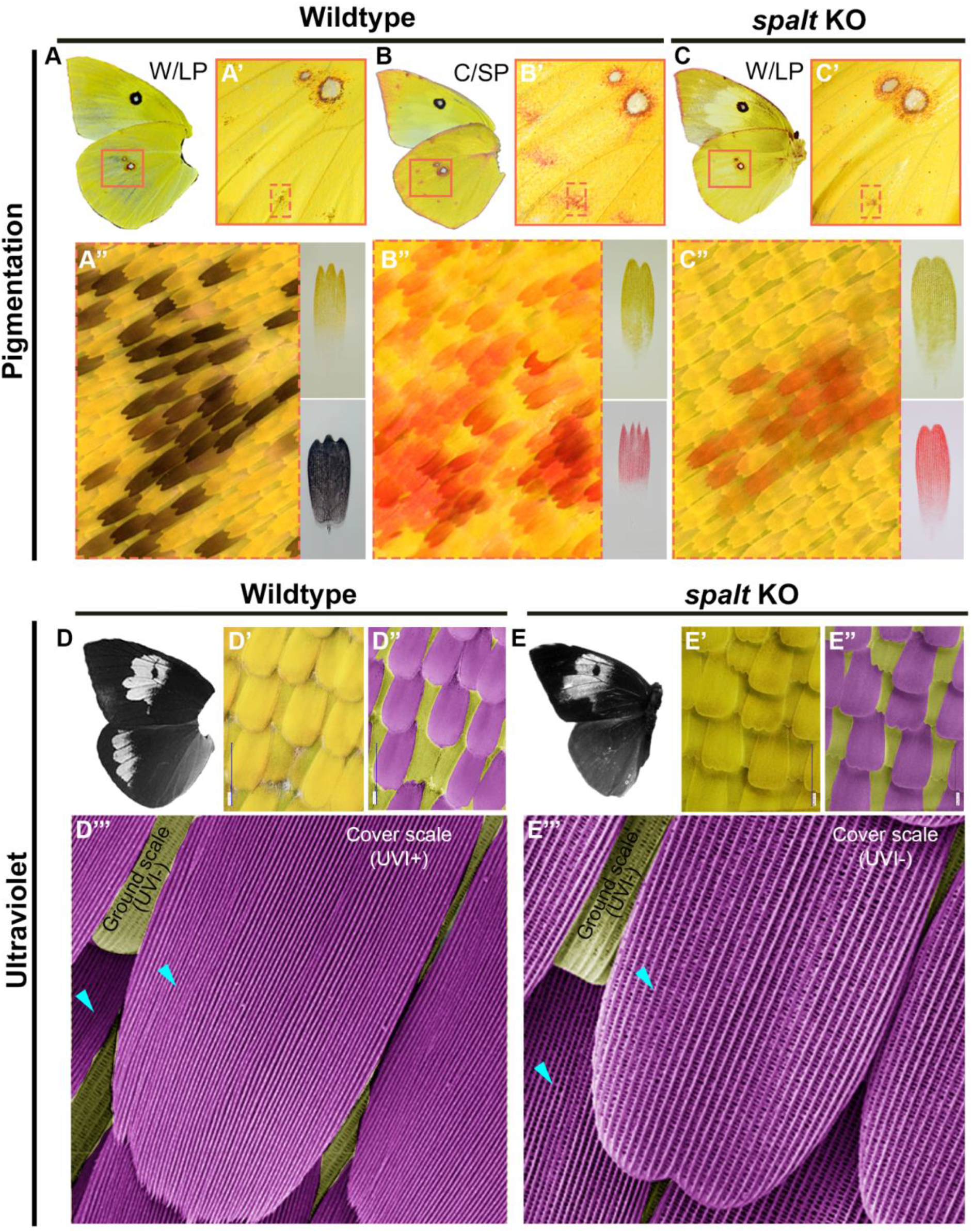
*spalt* KOs induce pink pigmentation and UVI plastic response. **A-A”** Melanic pattern and scales of ventral hindwings typical of summer and W/LP forms. **B-B”** Pink pattern and scales of ventral hindwings typical of winter and C/SP forms. **C-C”** *Spalt* crispants of W/LP reared butterfly results in pink pattern and scales typical of winter and C/SP forms. **D** UVI pattern typical of summer and W/LP forms. **D’** Light microscopy image of scales from UVI reflecting hindwing region in D. **D”** same image as D’, with non-UVI reflecting ground scales (yellow) and UVI-reflecting cover scales (violet) pseudo colored. **D”’** SEM of UVI reflecting hindwing region, with ground and cover scales pseudo colored. **E** UVI pattern of *spalt* crispant of W/LP reared butterfly showing lack of hindwing UVI, typical of winter and C/SP forms. **E’** Light microscopy image of scales from UVI loss hindwing region in E. **E”** same image as E’, with ground scales (yellow) and cover scales (violet) pseudo colored. **E”’** SEM of UVI reflecting hindwing region, with ground and cover scales pseudo colored. Blue arrows point to scale lamellar ridges highlighting the tight organization in D”’ and variably wide organization in E”’.

To test if the winter pink morphs were able to warm more quickly than summer yellow forms, we conducted warming experiments on preserved, wild-caught butterflies that varied in the amount of pink coloration on their wings. We observed a strong, significant positive correlation (r^2^= 0.52; p = 7.05×10^−8^) between the percent of pink on the wings and the rate of warming (Figure S4). In general, females showed greater amounts of pink across their wings than males, and the darkly pink-colored females showed the fastest rates of warming.

### Pink and UVI plasticity have a shared genetic architecture

During wing development in C*olias* and other Pierid butterflies, *spalt* is expressed where melanic discal cell and marginal spots form (13). We used CRISPR to test if *spalt* may be involved in the plastic change of melanic to pink in the hindwing discal cell and marginal spots of *Z. cesonia. spalt* crispants reared in W/LP conditions resulted in pink patterns in the discal cell and marginal spots, like those seen in C/SP reared butterflies (Figure 4). Detailed examination of the discal cell and marginal spots revealed that most scales were darkly pink pigmented. In addition, two crispant individuals had large white scales dispersed across the ventral hindwing, which are typically only found on wings of C/SP morphs (Figure S3-4)

**Figure 4.**
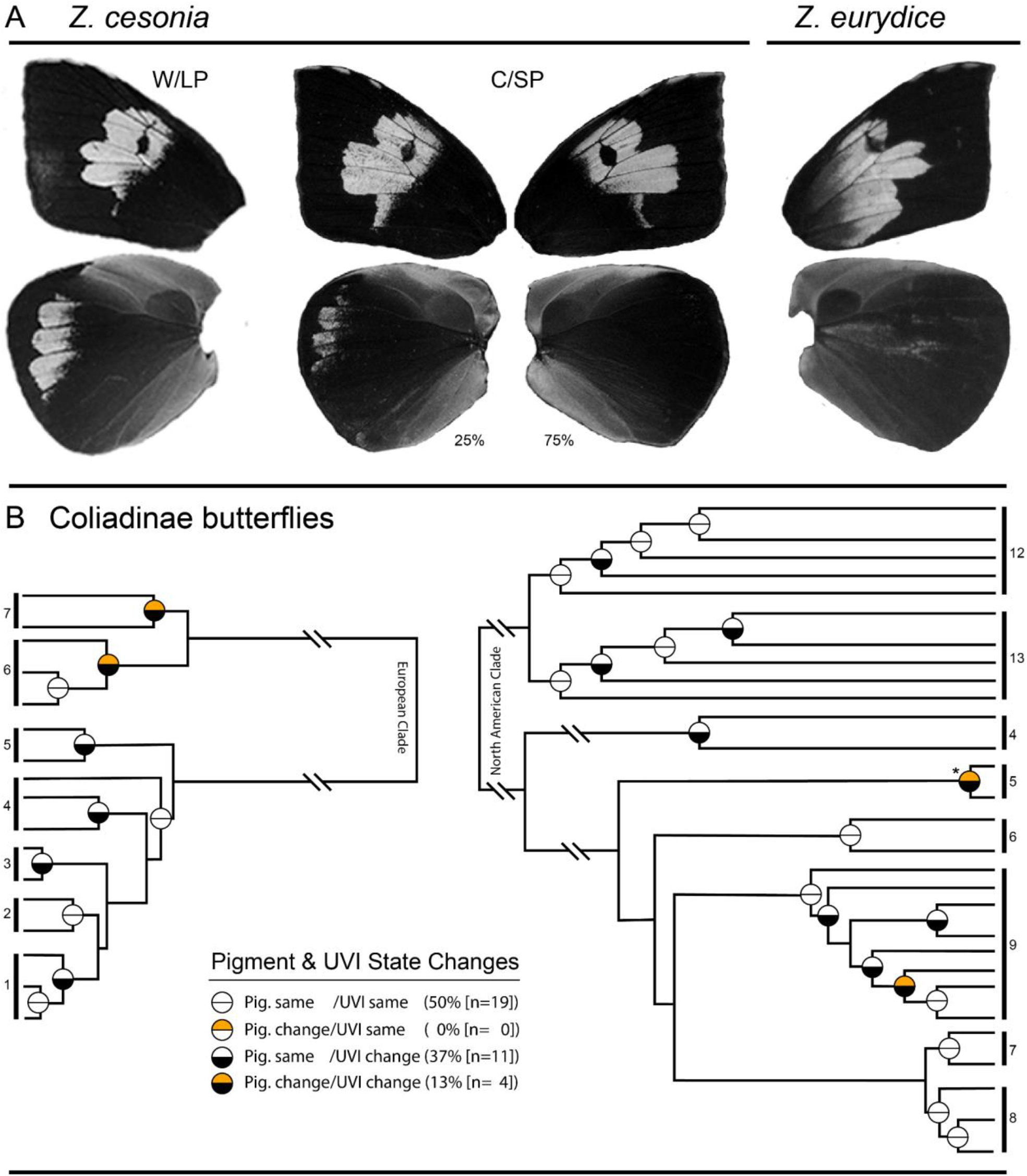
UVI plasticity phenocopies divergence between species. **A**. Cryptic genetic variation for plastic UVI loss in *Z. cesonia* phenocopies hindwing UVI divergence with *Z. eurydice*. **B**. UVI and pigment state changes among European and North American Coliadinae species groups (See Figure SXX). State changes reflect a difference in presence/absence of UVI or pigment pattern across hind, fore, or both wing dorsal surfaces.

Surprisingly, *spalt* editing also resulted in a complete loss of UVI on the dorsal hindwings in three of the seven male cripants (Figure S5). High magnification imaging of these scales lacking UV reflectance in *spalt* crispants revealed that the cover scales lacked the tightly organized lamellar ridges typically responsible for the UVI (Figure 4). However, there were notable differences between the cover and ground scale lamellar ridges, as well as noticeable variation in the distance between lamellar ridges of cover scales. This contrasts with non-UV reflecting yellow cover and ground scales on *Z. cesonia* wings, which typically have similar widely-spaced lamellar ridges, as well as the typical UV-reflecting cover scales that show very tightly organized lamellar ridges with little variation between scales (21). This variation in lamellar ridge organization among scales in this region may be due to mosaicism commonly observed on crispant butterfly wings (22, 23). Alternatively, this variation may be the result of *spalt* editing being insufficient to fully transition between UVI+ and UVI-scale ultrastructures.

It is noteworthy, that the *spalt* editing only impacted UVI and pigmentation in wing regions that show seasonal plasticity. Our results suggest that *spalt* may be involved in the plastic response of pigmentation and UVI in *Z. cesonia*.

### UVI plasticity phenocopies species divergence in Zerene and other sulphur butterflies

An interesting observation is that the absence of hindwing UVI in C/SP reared *Z. cesonia* males results in a hindwing that resembles the sister species, *Z. eurydice*. In this regard, the plasticity of UVI on the hindwing phenocopies the species differences in UVI on the hindwing. More importantly, only 75% of C/SP reared *Z. cesonia* males lacked UVI, suggesting there is genetic variation for the UVI plastic response. This variation in UVI plasticity could be interpreted as “cryptic genetic variation” since the different plastic responses are only observed when exposed to cold environments (note that this variation was observed in C/LP and C/SP treatments; see Figure 2). Among the 25% of C/SP individuals with UVI hindwing pattern present, morphometric analysis showed that the pattern clearly differed from W/LP and was more similar to wild-caught males from colder months (Figure S1D).). Like the observations of wild-caught males lacking UVI in cooler months, the C/SP reared individuals that lacked hindwing UVI all had >6% pink on the hindwing (Figure S1). Thus, in *Z. cesonia* there is genetic variation for the UVI presence/absence plastic response, the UVI plasticity segregates in a switch-like fashion, and this UVI plasticity phenocopies the UVI divergence with *Z. eurydice*.

Next, we wanted to determine if the UVI presence/absence differences on wing surfaces seen in the *Z. cesonia* plastic response was a common feature of species divergence among sulphur butterflies. To explore this, we examined UVI wing patterns of 46 European and North American Coliadinae species to determine if there were changes in the presence/absence of UVI on the dorsal surfaces of forewings, hindwings, or both wings between closely related species. We did the same for pigment (pterin and melanin) patterns, to compare the frequency of UVI and pigment divergence among sulphur butterfly species. Among the 30 groups of closely related species we studied, 50% showed changes in UVI, but only 13% showed pigment changes (Figure 5). Of the pigment changes, only one involved changes in melanin, and the other three cases involved orange pterin-based changes among *Gonepteryx* species (Figure 5, SYY). These results demonstrate that UVI presence/absence divergence similar to the UVI plasticity seen in *Z. cesonia* is frequently observed among closely species across the phylogeny of sulphur butterflies.

## Discussion

### Plasticity and diversification of Coliadinae butterflies

In this study, we integrate several critical pieces of evidence to assess the conditions required for plasticity in UVI to have facilitated the species diversification of sulphur butterflies. First, we build on previous evidence of plasticity in *Colias* and *Eurema* sulphur butterflies to demonstrate that similar wing color seasonal polyphenisms occur in *Z. cesonia*. We further show that the seasonal color variation has similar adaptive potential in *Z. cesonia*, with darker colored butterflies able to warm faster, and thus likely increasing their ability to mate and lay eggs in cold conditions. Importantly, we show some of the first evidence of concordant seasonal-induced plasticity in pigment and structurally-based coloration.

As demonstrated by Waddington, if a species varies in its plastic response, selection can act on that variation to drive divergence in the trait among populations. Interestingly, the genetic variation for UVI plasticity in *Z. cesonia* phenocopies the hindwing UVI differences with *Z. eurydice*. UVI patterns have repeatedly been shown to influence inter and intra-specific mate choice in sulphur butterflies (14–20). In *Colias*, UVI presence/absence differences between hybridizing *C. eurytheme* and *C. philodice* is the dominant isolating barrier (19, 20). Thus, there is ample evidence in sulphur butterflies for the potential of sexual selection to act on UVI variation. The genetic variation for plastic response of UVI in *Z. cesonia* creates an opportunity for natural or sexual selection to drive genetic assimilation that could influence adaptive divergence and speciation. Unfortunately, nothing is known about mate preference or isolating barriers between *Z. cesonia* and *Z. eurydice*, however, they have been reported to come into contact and hybridize in nature (24, 25). Exploring how UVI may influence intra-and interspecific mating in *Zerene* would offer critical insights into the potential of the UVI plasticity to have impacted speciation in these butterflies.

In sulphur butterflies, UVI has been shown to frequently be involved in mate choice and is likely a sexually selected trait that has influenced speciation. In a phylogenetic examination of UVI species differences across Coliadinae butterflies of Europe and North America (26, 27), We found that 50% of species pairs differed in UVI presence/absence on wing surfaces similar to the plastic variation seen in *Z. cesonia*. Thus, supporting that the UVI color pattern differences generated by plasticity are similar to the UVI differences commonly found between closely related Coliadinae species. These findings highlight the potential of seasonal plasticity to phenocopy and influence the diversification of sulphur butterflies.

The prevalence of UVI divergence among closely related sulphur species supports UVI being a sexually selected trait that could have influenced species diversification (14). UVI is often sexually dimorphic and can vary dramatically among sulphur butterfly species (28, 29). There is support for this UVI diversity having a sexually selected origin using a similar phylogenetic approach that we employed (14). This earlier study quantified UVI and melanic patterns among nine species groups of sulphur butterflies and found support for a polyphyletic distribution of UVI patterns, indicative of a sexually selected trait. Similar to our findings in *Z. cesonia*, that plasticity responses are independent between the hind and forewings (e.g. UV only changes on hindwing), divergence among sulphur species also was independent among hind and forewings (Figure 4B, S6-7). These findings support that the UVI variation generated by plasticity is similar to the UVI differences that are commonly found and known to influence sexual selection among closely related Coliadinae species.

Together, our findings offer a string of evidence that supports the potential for plasticity in UVI to have influenced the diversification of sulphur butterflies. However, there are several key pieces of evidence that remain to be tested. Perhaps the most critical is evidence that genetic assimilation of UVI can and has occurred. Studies of seasonal polyphenism in *Junonia* suggest that genetic assimilation of butterfly wing color plasticity can occur in natural populations (12). Demonstrations that it has occurred in the evolutionary history of a species are much more difficult to discern and in sulphur butterflies, will inquire further investigation of the ubiquitousness of pigment and UVI wing color plasticity. Dissecting the genetic architecture of genetic assimilation would also be a useful approach, as we could then explore the divergence and function of genes involved in genetic assimilation among natural populations of sulphur butterfly species. Regardless, we provide compelling evidence that there is plasticity of UVI in *Z. cesonia* that could influence species divergence and highlight seasonal plasticity of wing colors in sulphur butterflies as a promising system to explore the impact of developmental plasticity in adaptation and speciation.

### Pigment and structural color plasticity are linked by the environment and genetics

The shared similar environmental cues for pterin pigmentation and structurally-based UVI plasticity in our study suggest plasticity in UVI could be as common as pigment plasticity in sulphur butterflies. Plasticity of pigment-based wing color patterns in response to changes in temperature and/or photoperiod is a hallmark of many butterfly species (3). Using museum collections, MacLean et al (30) demonstrated the effects of climate change on the plasticity of wing melanization in *Colias meadii*. Using a similar collections-based approach, we surveyed wing color among image collections of sulphur species (28, 29) and found variation in black, pink and/or UV wing patterns that are similar to the plastic wing patterns of *Colias* and *Zerene* among species from almost every genus of Coliadinae butterflies (13 of 17 genera; see Supplemental Material; Figure S8). Thus, seasonal plasticity of pigmentation patterns is likely common among sulphur butterfly species.

Despite the commonness of pigmentation plasticity on butterfly wings, there are only a few examples of plasticity in structurally-based colors. In *Pieris rapae*, a close relative to sulphur butterflies, non-iridescent UV has been shown to vary seasonally (31). In *P. rapae*, UV variation is a direct result of the density of pterin pigment granules, which are highly UV absorbent, rather than changes in any UV-reflecting nanostructures typical of Coliadinae butterflies. In the distantly related lycaenid butterfly, *Polyommatus icarus*, cold stress was shown to impact both pigment and structurally-based blue wing coloration (32). Our study and the others consistently show that coordinated changes can occur in pigment with UV and/or other structural colors in response to environmental cues.

The genetic coupling of pigment and UVI changes as seen in our *spalt* KOs offers insight into the mechanisms that may be driving the coordinated plastic responses of pigmentation and UVI. In the *Z. cesonia spalt* KOs pink and UV changes occurred broadly across different wing surfaces and regions. This contrasts with *spalt* expression in *Colias eurytheme* that was highly localized to the melanic regions of the wing (13). This suggests that the plastic response involves an environmentally sensitive signal that is broadly distributed across the wing and upstream of *spalt*. A similar model was proposed to explain eyespot plasticity on the wings of *Bicyclus anynana*, where environmentally-induced changes in wing titers of 20-hydroxyecdysone (20E) are associated with localized expression changes of Ecdysone Receptor (EcR) in the eyespot centers (33). Further study of the spatial expression of *spalt* and the influence of environmental changes of hormones, such as Juvenile Hormone and 20E across developing wings of *Z. cesonia* should be explored to elucidate the developmental mechanisms involved in generating pigment and structural color plasticity.

## Methods

### Butterfly collection

Adult *Z. cesonia* were collected from February through October from Osborn prairie (33°30’36.98”N, 88°44’14.57”W) in 2016. In the laboratory, both sexes were released into flight cages, allowed to mate, and females were allowed to oviposit on *Dalea purpurea*. After death, the wings of both sexes were preserved to track the phenotypic changes in wing color patterns throughout the season.

### Larval rearing and treatment conditions

Eggs laid from wild-caught females in May 2016 were collected daily and placed into a designated environmental treatment chamber. After hatching, larvae were reared on an artificial diet, and after eclosion, 30 individuals from each treatment were sacrificed and preserved for imaging. The eggs were split among rearing conditions Warm/Long Photoperiod (W/LP; 27°C/16:8 hrs), Warm/ Short Photoperiod (W/SP, 27°C/8:16 hrs), Cold/Long Photoperiod (C/LP; 21°C/16:8 hrs), and Cold/Short Photoperiod (C/SPi; 21°C/8:16 hrs), and an additional C/SP (C/SP; 25°C for 10hr; 16°C for 6hr; 10°C for 2hr; 16°C for 6hr/10:14 hrs). These conditions were chosen to best match temperature and photoperiod conditions from the local region using data acquired from the National Ocean and Atmospheric Administration weather database (https://www.ncdc.noaa.gov/) for weather station GHCND:USC00228374 (33.4691°, - 88.7822°). The initial C/SPi rearing with the 21°C/8:16 hrs C/SP conditions induced a high incidence of diapause, preventing eclosion and study of wing color patterns. The latter C/SP conditions did not induce diapause responses, and better reflected the nighttime low temperatures experienced in local populations during March and October 2016, as was done for the Warm conditions.

### Microscopy and Imaging

Wings were photographed using a Nikon D7000 camera with an AF-S Micro Nikkor 105 mm Lens for visual color images. Ultraviolet color images used the same lens and camera but with the addition of a BAADER U-Filter, 60 nm IR and VIS filter. Light microscopy images of scales on the wing were taken on a Leica M205C dissecting scope with a Cannon 650D camera or a Keyence VHX-7100. Images of individual scales removed from the wing were taken with light microscopy on an Axioskope 2 plus (Carl Zeiss) microscope with a Cannon 5DS camera or a Keyence VHX-7100. Scales were removed from the wing using a tungsten needle and submerged in clove oil, which has a similar refractive index as insect cuticle (Wasik et al, 2015), thereby limiting the structural color reflectance. SEM images were collected using the methods described in Fenner et al 2019 (21).

### Color pattern analyses

Pink and UV across the hindwing surfaces were quantified by transforming color images. Using Fiji (34), color and UV wing images were transformed to binary black and white images. In color images pink elements were transformed to black on a white background, and in UV images the UV elements were transformed to white on a black background. Calibrated areas were measured for the pink elements, UV elements and total hindwing size. The percent pink and UV values were standardized by the total wing area. T-tests were performed in Rstudio to test for significant differences in percent pink and UV among sample groups.

Additional landmark-based geometric morphometrics were also performed to quantitatively measure changes in the UV patterns between the laboratory and wild males. Twelve landmarks were chosen and placed on the hindwing to encompass the entire UV pattern. Landmarks consist of Type I, placed at wing veins, Type II, placed at the maximum points of the UV pattern, and Type III, anterior and posterior points of the structure (35) using the ImageJ package Pointpicker (36) and exported to MorphoJ (37). A procrustes fit, aligning by the principle axis, was performed on the data set and a covariance matrix was produced using the procrustes coordinates. Canonical variant analysis was performed from the covariance matrix and plotted in a scatterplot with 95% confidence ellipses.

### Warming experiment

The rate of warming was conducted on 40 preserved butterflies that ranged from completely yellow (0% pink) to maximally pink (80% of the hindwing was pink). Butterflies from the field and laboratory experiments were preserved in glassine envelopes and allowed to desiccate for 16-24 months. Individual butterflies were placed in a black 12-inch cube chamber, made of one-inch thick black foam. The cube chamber had no top, allowing for an overhead heat lamp. The bottom of the cube chamber was lined with white poster board. Eight thermocouples were inserted through the foam, and the wires were taped to the white poster board, approximately one inch from the end of each thermocouple. To calibrate before each set of readings, the chamber was cooled and maintained at 25°C for three minutes. After three minutes, the heat lamp was turned on, and the ambient temperatures in the chamber were recorded. The setup was then calibrated by adjusting the heat lamp position and distance to achieve an increase from 25 to 37°C in 3 minutes. This calibration was chosen to match the temperature range and rate of warming observed in Colias butterflies. Once the setup was calibrated, sets of six preserved butterflies were placed on the white poster board with their right ventral wing on the surface. Thermocouple probes were inserted between the closed wings, firmly touching the dorsal side of the thorax. Two probes were left without butterflies to record the ambient warming rate. This was replicated three times for each group of six butterflies, with the positions of the butterflies switched between experiments. The rate of warming was estimated by taking the ratio of a butterfly’s warming rate (seconds/14C) to the ambient warming rate (seconds/14C), and then averaging that across the three replicates. This ratio allowed us to control for variation in ambient environmental conditions in the laboratory between experiments. For each individual, the average rate of warming ratio and % pink on the hindwing were plotted, followed by a linear regression and a T-test to test the significance of regression.

### CRISPR-based gene editing

Single Guide RNAs (sgRNAs) were designed from a *spalt* transcript sequence from the *Z. cesonia* transcriptome (38). Guides were designed by manually scanning for areas of the transcript that are composed of 18-22 base pairs beginning with G and ending with NGG. Guide sequence was ordered from Integrated DNA Technologies as Alt-R® crRNA in Alt-R® CRISPR-cas9 system. The guide sequence was 5’-GCAAATGTTTTGTAGCAGATGG-3’. Guide complexes were formed by mixing Alt-R® crRNA with Alt-R® tracerRNA to form the guideRNA (gRNA). Alt-R® S.p. Cas9 Nuclease V3 and the sgRNA were combined and injected into developing embryos. Embryo injections occurred between 3-5 hours after eggs were laid and injected with a ∼1 ul of a 10 ul mix composed of 1ug of Cas9 and 100 ng of gRNA. Borosilicate glass needles were used for injections and eggs were affixed to glass slides with double-sided sticky tape. After injections, eggs were kept in a petri dish in a rearing chamber with 50% humidity, 16:8 hour photoperiod, and 27°C temperature (W/LP). After three days, host plant cuttings were laid across the slide allowing larvae to immediately climb onto host plants once hatching occurred on day four. Larvae were reared on unlimited *D. purpurea* till pupation. Upon emergence, adults were frozen, pinned, and either leg or wing snips were removed for genotyping. DNA was extracted with a Qiagen DNeasy Kit from legs flash-frozen in liquid nitrogen. Wing snips were immersed overnight at room temperature in 20ul TE and homogenized with a pestle. PCR using Q5 high-fidelity Taq polymerase was conducted using primers that flank the guide sequence. Forward primer 5’-GAACGGACGTTCTTTGGTGT-3’ and reverse primer 5’-GAAGGCGATGGTGGTGAG-3’. PCRs were cloned using pGEM®-T Easy Vector System and sequenced with capillary Sanger sequencing using BigDye Terminators.

### Phylogenetic color pattern analyses

We compared images of wing patterns for 46 Coliadinae species to determine if UV and pigment patterns varied in a similar fashion between species as we observed in the plastic response. UVI wing patterns were for each species were characterized as (0) no UV on forewing (fw) and hindwing (hw); (1) UV on forewing (fw), no UV on hindwing (hw); (2) no UV on forewing (fw), UV on hindwing (hw); (3) UV on forewing (fw) and hindwing (hw). Pink pigmentation was scored the same. Pigment patterns were similarly characterized. Phylogenies of European and North American clades of Coliadninae butterflies based on multigenic phylogenetic analyses were used to assign the species to pairs, or groups of closely related species (26, 27). For each species pair/group, ancestral states were reconstructed at each split using parsimony algorithms from the castor package (39) in the R programming language. For each species group, we then tabulated if there was a change in UVI and/or pigment pattern presence/absence and calculated the likelihoods of the ancestral state. Theoretically, the phylogenetic signal of a trait under sexual selection (UVI) is weaker than a trait not under sexual selection (pink pigmentation) (40). Alternatively, a monophyletic pattern of color pattern changes would support that color pattern is not likely a target of sexual selection or influencing speciation. This approach to testing for evidence of sexual selection acting on UVI is analogous to earlier analyses on this butterfly clade by Kemp et al. (14).

## Acknowledgments

We would like to thank the Friends of the Black Belt Prairie for access to collection sites, John Riggins for access to equipment for temperature assays, Mike Brown for providing temperature data, and Michael Miller at the Auburn University Instrumentation Facility for assistance with scanning electron microscopy images. We thank Arnaud Martin and members of the Counterman lab for thoughtful discussions and feedback on earlier versions of the manuscript.

## Author Contributions

BAC and JLF conceived and designed the study. JLF performed experiments. JLF, HS, and BAC performed butterfly collections. JLF, HS, and AC conducted imaging of butterfly wings. KE performed temperature experiments. VF conducted imaging and phylogenetic reconstructions. BAC, JLF, VF, and AC conducted data analyses. RR provided access to equipment and guidance for CRISPR experiments. BAC and JLF wrote the paper with input from all co-authors.

## Supplemental Material

**Figure S1.**
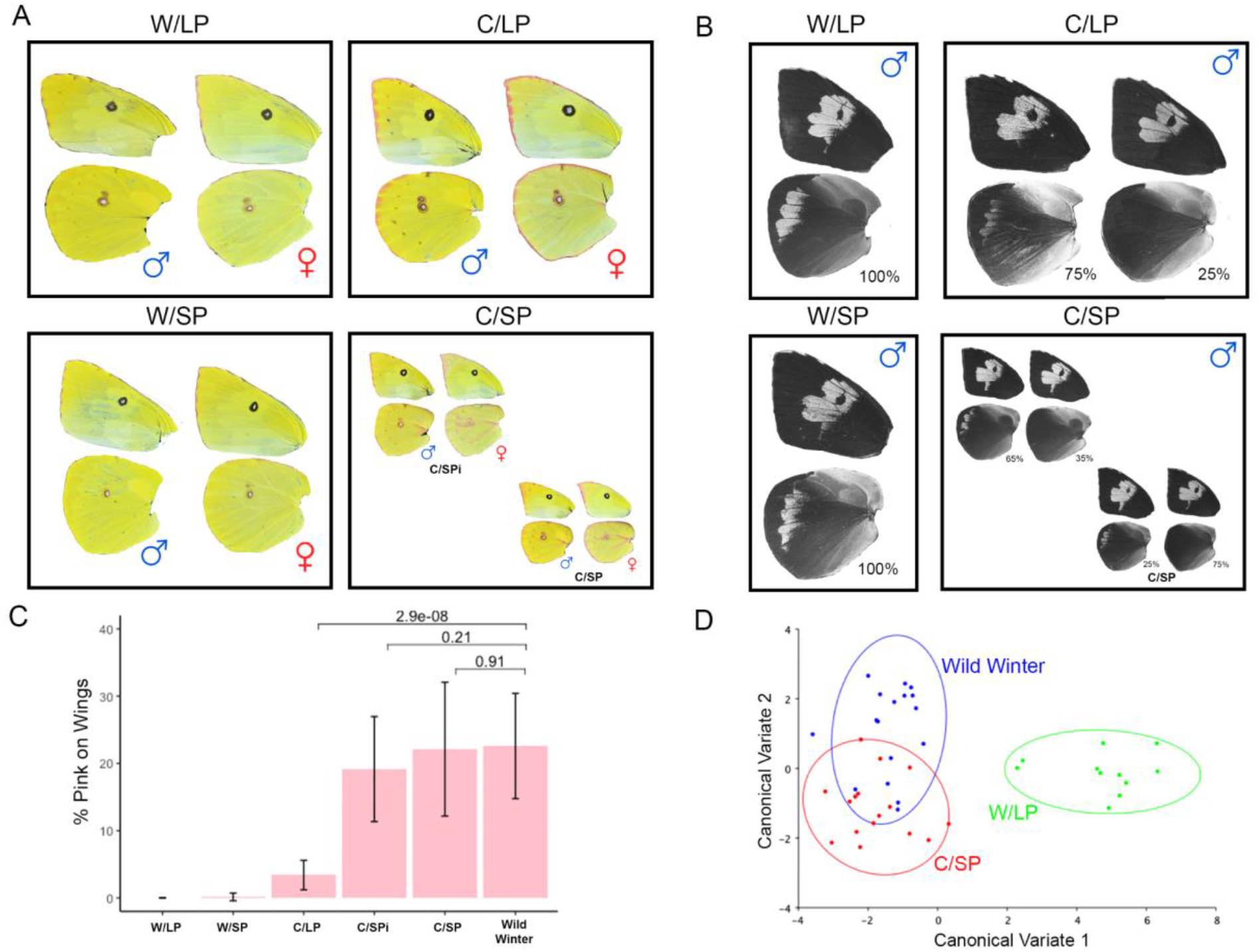
Wing pattern plasticity of *Z. cesonia*. **A-B**. Examples of male and female wing patterns in varying laboratory-controlled rearing conditions. **C**. Analyses of pink% of wings from varying rearing conditions. **D**. Canonical analyses of morphometric measurements of hind wing UV pattern among three rearing conditions.

**Figure S2.**
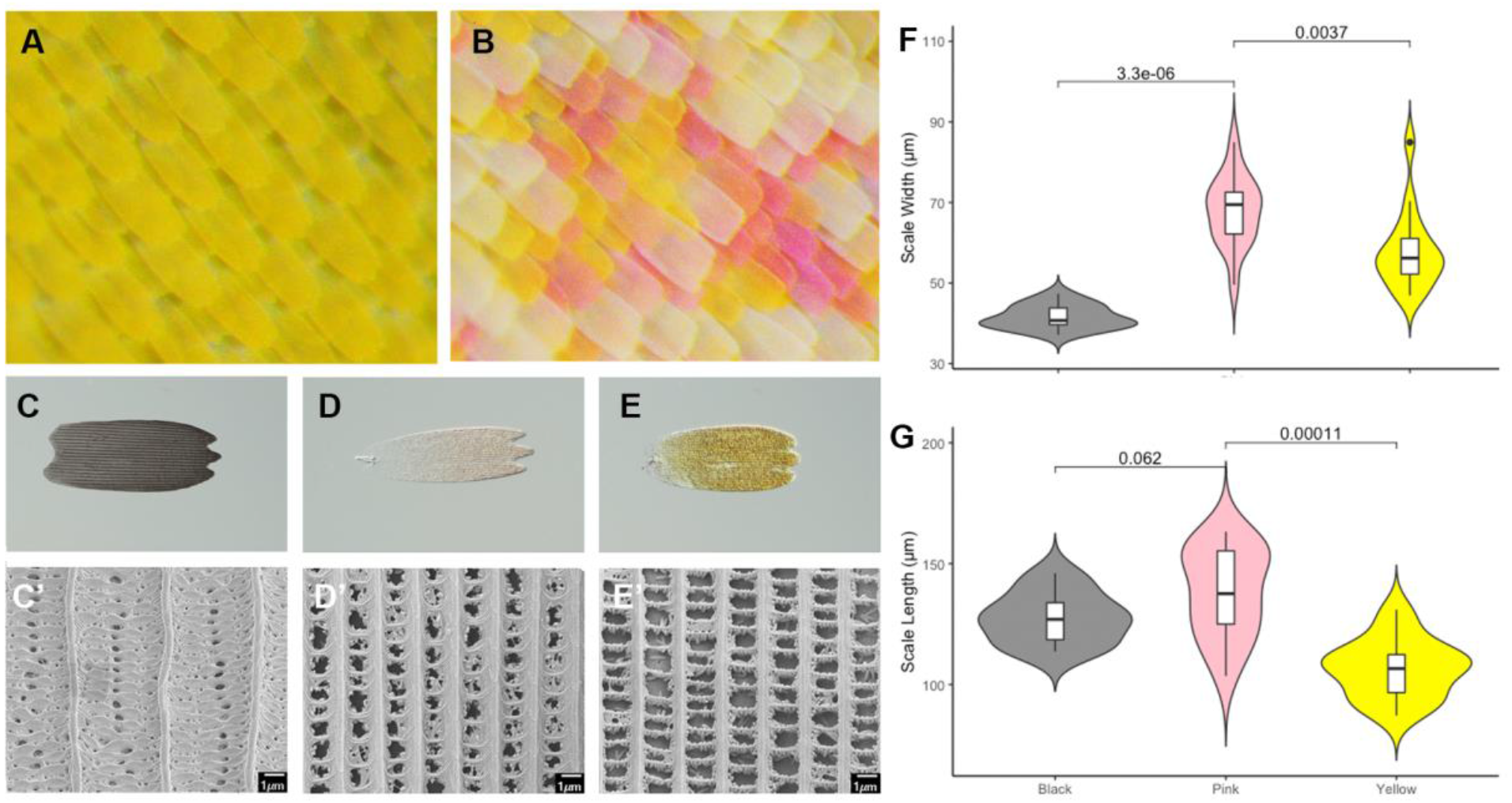
Scale diversity of summer and winter forms. **A**. Yellow scales typical of summer forms. B. Mix of yellow, pink and white scales typical of winter pink forms. **C-C’**. Melanic scale ultrastructure and lamellar surface typical of summer morphs. **D-E’**. Scale ultrastructure and lamellar surfaces of pink and yellow scales typical of summer morphs. **F-G**. Differences in scale width and length among summer form melanic scales and winter form yellow and pink scales sampled from homologous wing regions.

**Figure S3.**
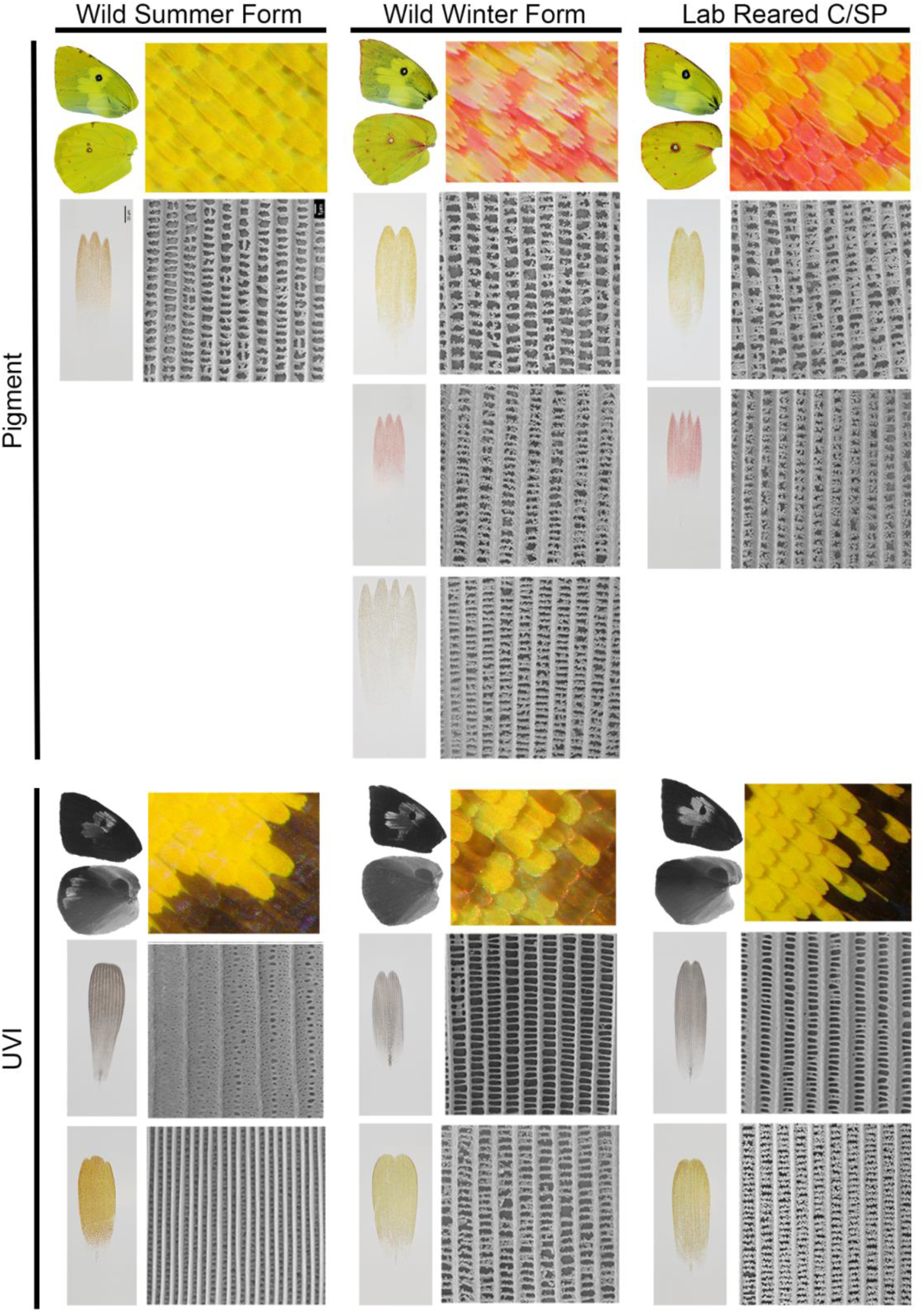
Examples of scale ultrastructure and lamellar surfaces for wild summer and winter forms, with lab-reared C/SP form.

**Figure S4.**
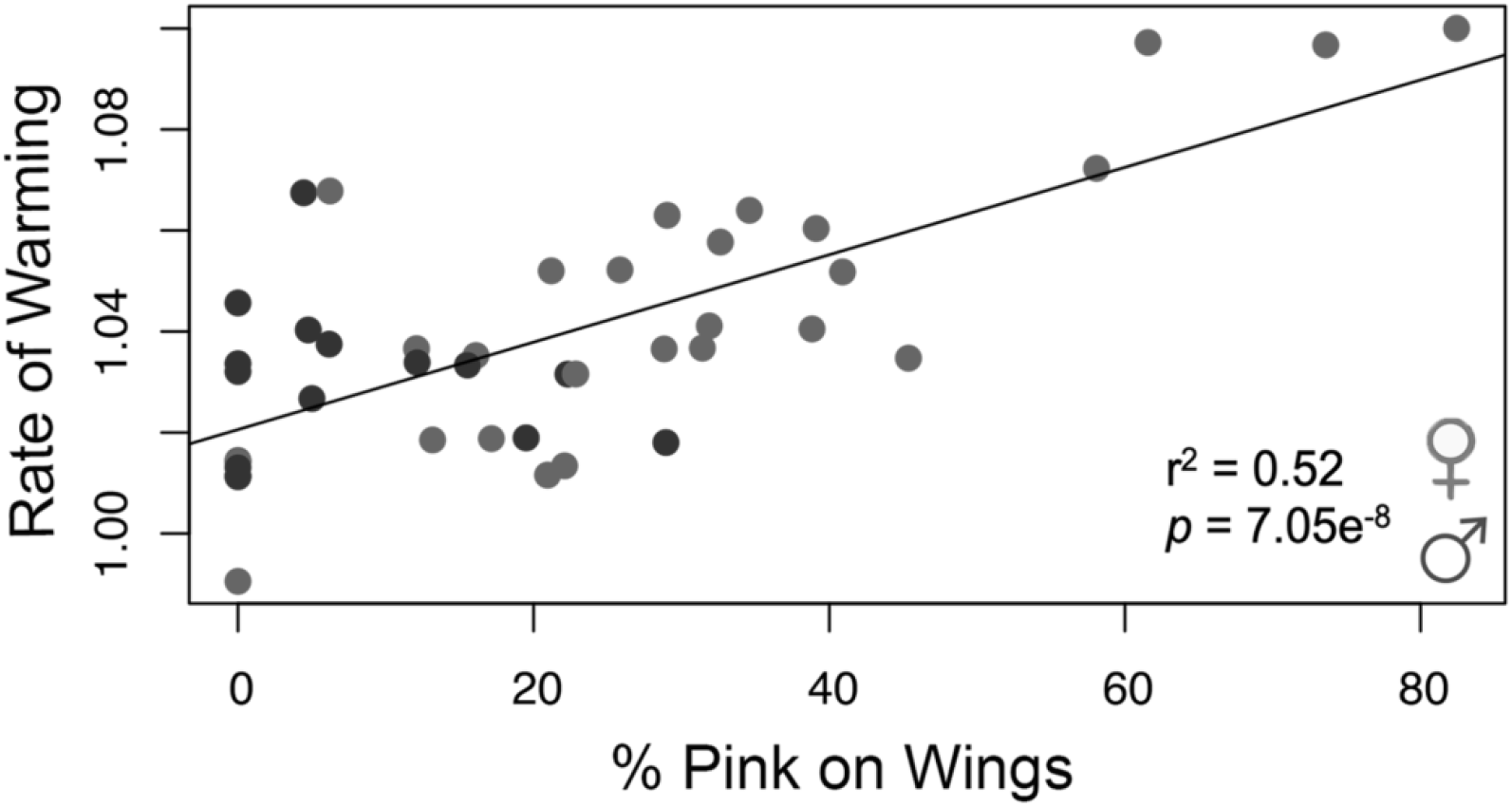
Rate of warming increases with pink pigmentation.

**Figure S5.**
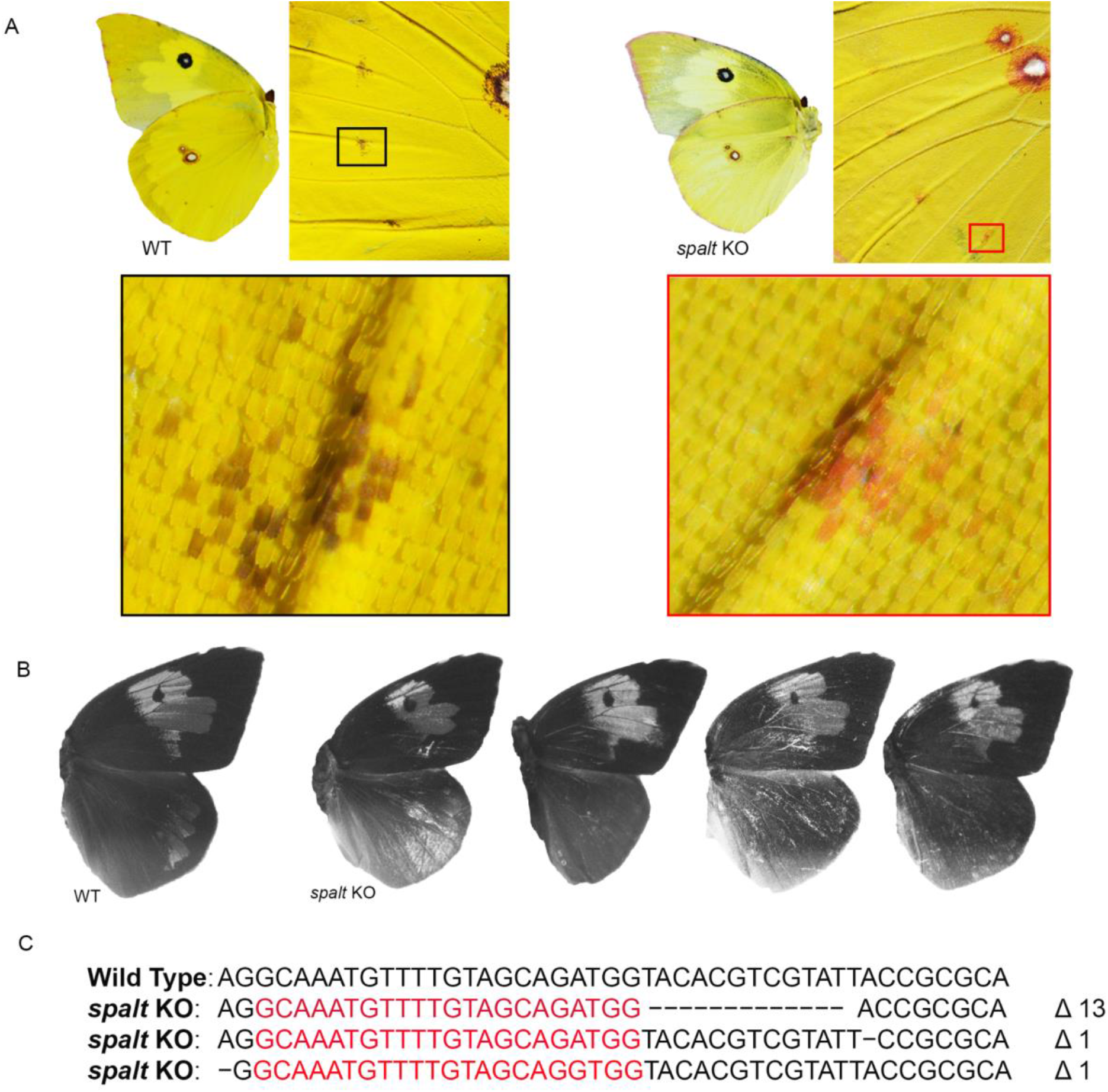
Additional *spalt* KOs. A. Wildtype and *spalt* crispant pigment patterns and scale types. **B**. Wildtype and *spalt* crispant UVI patterns. **C**. Examples of CRISPR edits to *spalt* sequence in KOs.

**Figure S6.**
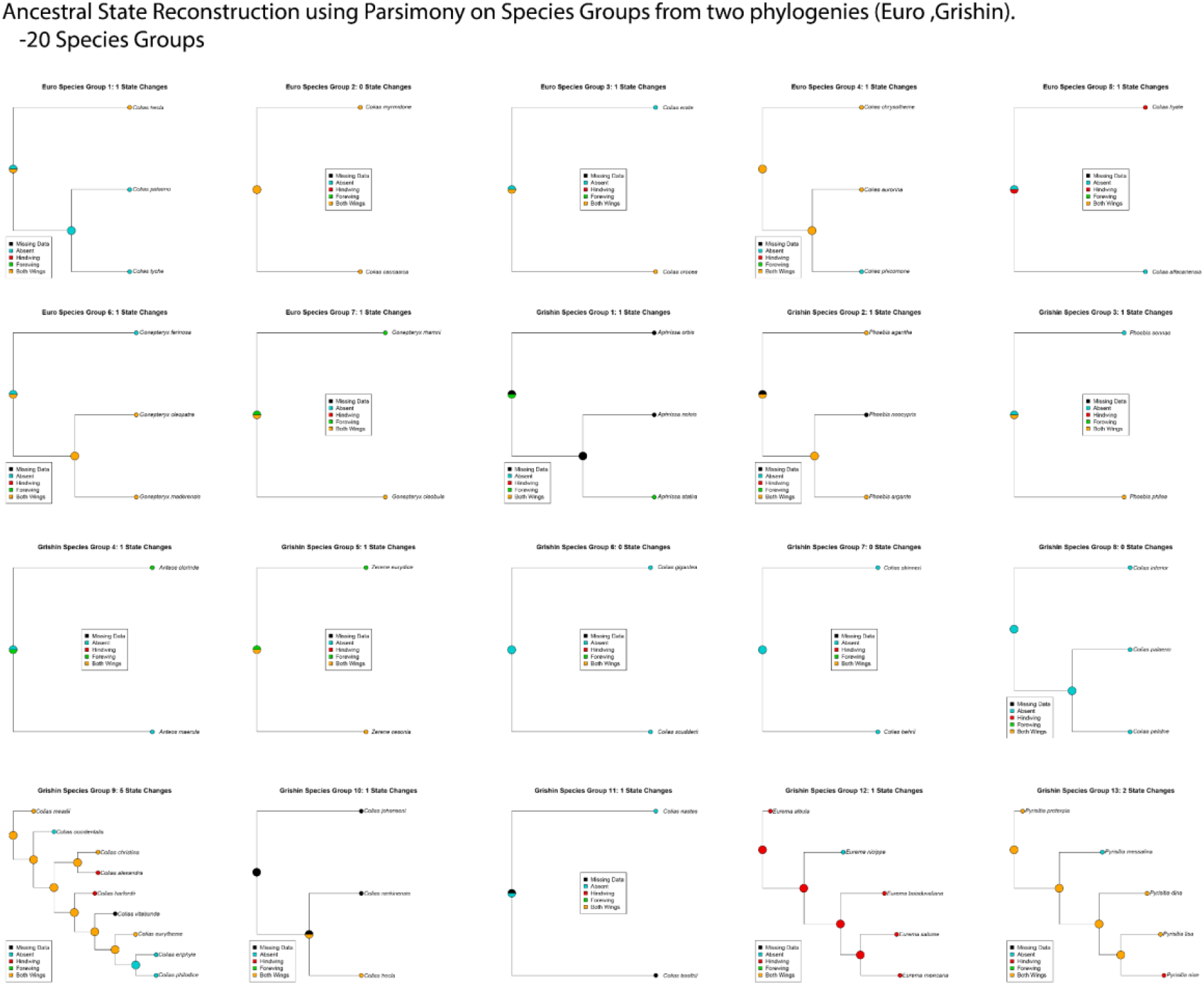
UVI state reconstructions for European and North American clades of Coliadinae butterflies.

**Figure S7.**
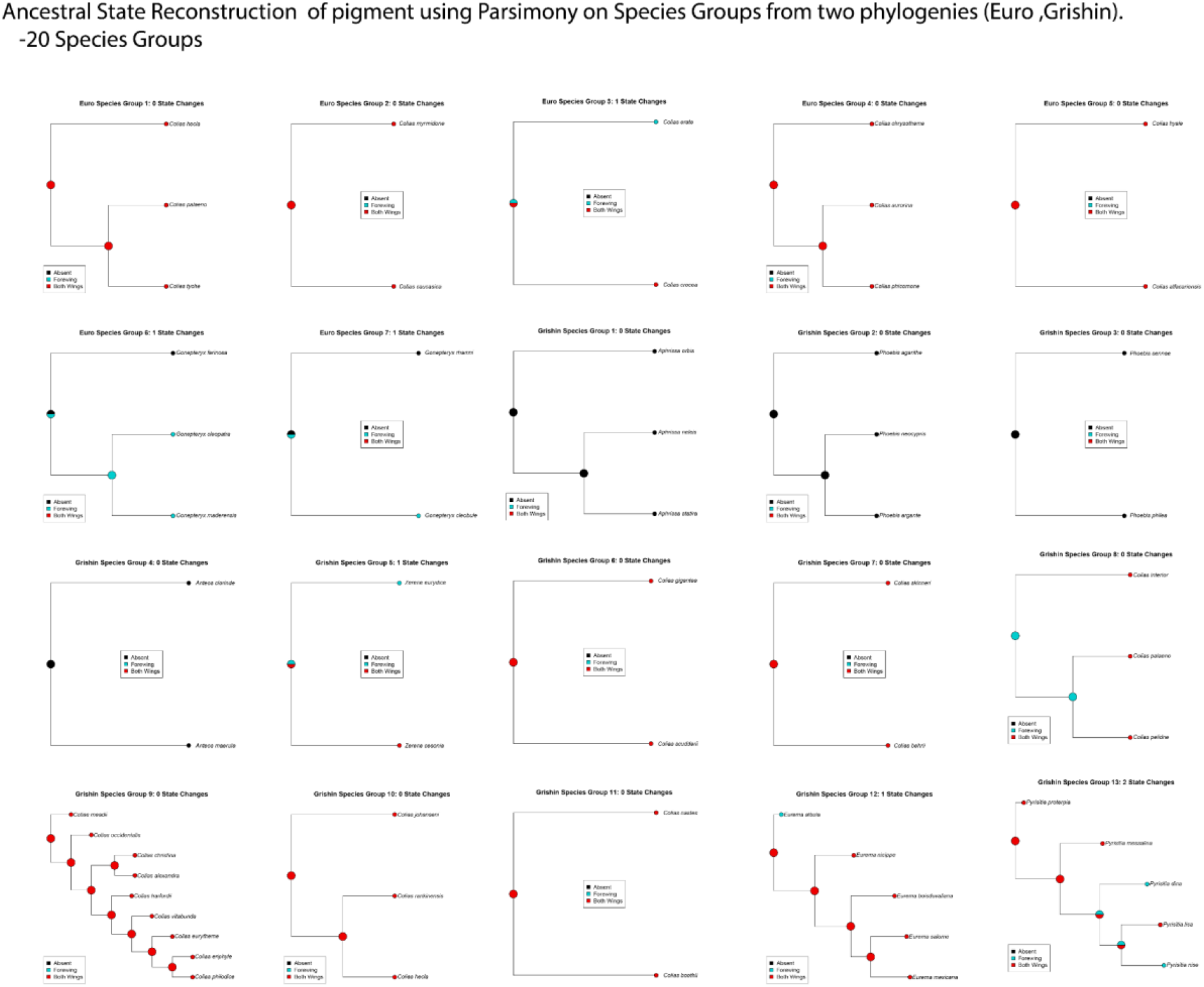
Pigment state reconstructions for European and North American clades of Coliadinae butterflies.

## Survey of wing color variation among Coliadinae

Plasticity of wing coloration has been described for several Coliadinae species, however, there have been no attempts to assess its prevalence across the clade. Here, we provide a survey of wing color variation among Coliadinae species from the scientific literature and collection images available through northamericabutteflies.org. We focus on addressing the questions: (1) is there evidence of wing color variation within species, and (2) does wing color vary in regions and colors in a similar fashion to the plastic responses in *Zerene* and *Colias*? For question 2, we used available images, reports from the literature, and/or images from butterfliesofamerica.com to confirm if pink or black coloration varied on the ventral FW tip, dorsal FW tip, or dorsal spots among collection images. For many Coliadinae species, this online database includes the type specimen for the species, as well as several other individuals of both sexes to assess the presence of variation.

Four of the 17 Coliadinae genera surveyed (*Anteos, Luecidia, Gandaca* and *Dercas*), had insufficient numbers of individuals to make assertions of wing color variation (Figure S8). Among the other genera, all showed evidence of wing color pattern variation within species that resembled the seasonal plasticity of *Zerene* or *Colias*.

**Figure S8.**
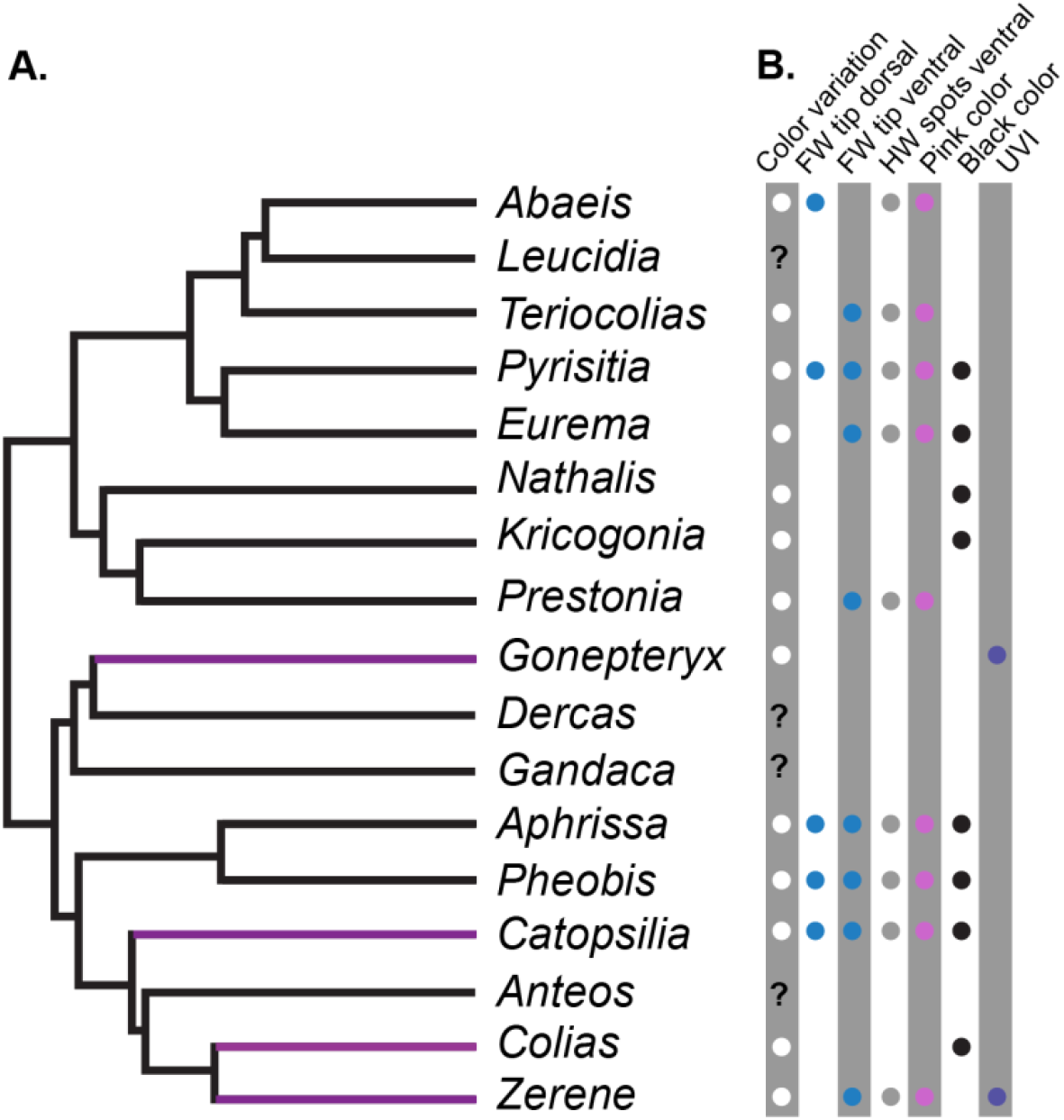
Survey of wing color variation among Coliadinae genera. **A**. Phylogenetic relationships of Coliadinae genera taken from (41). Purple branch mark genera with species that have published evidence of wing color plasticity in response to temperature, photoperiod, and/or precipitation. **B**. Description of wing color variation observed within species. Question marks reflect insufficient data in collections to make assertions of wing color variation. Color variation = is there evidence of wing color variation in collections; FW dorsal tip = is there change in the thickness of melanic region on FW dorsal distal edge; FW ventral tip = is there change in the thickness of melanic region on FW ventral distal edge; HW ventral spots = are there changes in pigmentation and/or size of discal and marginal wing spots on ventral surface; pink = do color changes involve pink coloration; black = do color changes involve black coloration.

*Nathalia iode* and *Kricogonia lyside* show color variation similar to *Colias* species. In *Colias*, seasonal polyphenism of wing coloration has been well characterized and is known to occur in response to changes in photoperiod in juvenile stages in *Colias eurytheme* (10). Darker morphs, marked by increased melanic scale coverage across the wing, are more common at cold temperatures. The darker coloration allows the butterflies to warm/cool more rapidly (9), thereby increasing flight times and the opportunity for mating and oviposition. The degree of melanin across the wing varies within and between many Colias species. The increase of melanin appears to be the result of melanin scales speckled across the typically uniform yellow or white wing surfaces. The greatest increase in melanin appears in the proximal wing regions nearest the thorax, particularly on the ventral wing surface. This type of increase in melanin across the wing is seen in the collections of several *Colias* species.

*Eurema, Abeis, Pyrisitia, Teriocolias* and *Prestonia* all had multiple species that showed interspecific variation in wing regions and colors that vary in *Z. cesonia* plastic responses to temperature and photoperiod. Interestingly, all these genera have species that show similar patterns of variation in pink pigmentation. The changes in pink pigmentation are most obvious on the ventral hindwing surface and the distal tip of the ventral forewing surface. The hindwing color variation are most obvious at marginal and discal spots, but can also extend broadly across the forewing, very similar to observations of plastic wing colors in *Z. cesonia*.

*Phoebis sennae* showed an increase in melanin on dorsal wing margins and an increase of pink coloration on ventral surfaces of fore-and hindwings. Similar to observations in the genera above (e.g. *Eurema* etc.), *Phoebis* intraspecific wing color variation largely involves enlargement of spots across wing surface. Unlike species from the genera above, *Phoebis sennae* color variation occurs broadly across forewing and hindwing ventral surfaces.

*Catopsila* (42) *and Aphrissa* show intraspecific variation in melanin on the dorsal forewing margin, which is not observed in *Z. cesonia* plastic colorations, but there is evidence that the melanic margin may change in size in plastic responses of *Colias* species. However, variation in this region of the forewing has not been quantitatively assessed or formally described as a plastic response to temperature or photoperiod, to our knowledge. Species from these genera also show suggestive evidence of variation in pink coloration on the dorsal surfaces, particularly at discal spots, marginal spots, and the distal forewing tip.

*Gonepteryx rhamni* shows clear variation in ultraviolet patterns in response to the developmental environment (28), but no obvious variation in pigmentation. Pechacek et al (28) showed that the majority (93%) of quantitative variation in UV pattern among Palearctic *Gonepteryx rhamni* could be explained b variation in precipitation, temperature and latitude. This is the only species other than *Z. cesonia* that UV plasticity has been assayed.

